# 5-HT2A receptor deletion is associated with reduced hippocampal parvalbumin levels and anxiety-like behavior in BALB/c mice

**DOI:** 10.64898/2026.11.19.767898

**Authors:** Katelyn Dial, Pauline Wonnenberg, Junfeng Ma, Katherine Conant

**Affiliations:** Department of Biochemistry and Molecular & Cellular Biology, Georgetown University, Washington, DC, United States of America; Department of Neuroscience, Georgetown University, Washington, DC, United States of America; Mass Spectrometry and Analytical Pharmacology Shared Resource, Georgetown University, Washington, DC, United States of America

## Abstract

The serotonin 2A receptor (5-HT2AR) is a target of drugs used to treat anxiety and mood disorders, yet whether endogenous 5-HT2AR signaling constrains or promotes anxiety, and how the 5-HT2AR affects cell populations linked to anxiety, is not fully resolved. Acute 5-HT2AR agonism is anxiolytic and acts through the activation of ventral hippocampal parvalbumin-expressing interneurons. Here we report that constitutive deletion of the 5-HT2AR in male and female BALB/c mice is associated with robust anxiety-like and repetitive behavior across multiple paradigms. Unbiased proteomic analysis of male knockout hippocampal tissue revealed substantially reduced levels of parvalbumin (PV), a proxy of their activity. Consistent with the intense metabolic demands of fast spiking interneurons, proteins supporting energy metabolism were also downregulated, including those related to oxidative phosphorylation and the electron transport chain, glycolysis and the tricarboxylic acid cycle, together with the cytoplasmic ribosome. In terms of how reduced 5-HT2AR signaling could lead to reduced PV expression, we examined key perineuronal net (PNN) components. The PNN constituents aggrecan and HAPLN1 were not significantly changed, whereas the link protein, HAPLN4 was substantially reduced. In a separate cohort, 7 days following a single dose of the selective 5-HT2AR agonist, 25CN-NBOH, many of the same energy metabolism proteins were shifted in the opposite direction. Together these findings indicate that constitutive loss of 5-HT2AR signaling is associated with coordinated reductions in parvalbumin and in the biosynthetic and bioenergetic machinery that supports fast-spiking activity, without a generalized loss of perineuronal net structural components.

## Introduction

The serotonin 2A receptor (5-HT2AR) is expressed at high levels in cortical and limbic brain regions including prefrontal cortex, amygdala, and hippocampus [1–3], where it is the most abundant excitatory serotonin receptor and couples canonically to Gq [4]. It is a target of drugs used to treat anxiety and mood disorders, from the atypical antipsychotics to the psychedelic compounds now in clinical trials [5–7]. Work on how the receptor shapes emotional behavior has concentrated largely on cortical pyramidal neurons, where the 5-HT2AR is enriched on apical dendrites and receptor activation increases pyramidal excitability [8], and where the behavioral consequences of receptor knockout and the downstream effects of agonists have been well studied [9,10]. Within the hippocampus, however, the consequences of 5-HT2AR knockout have been less well explored.

Recent work has moved hippocampal inhibitory interneurons to the center of this question. The 5-HT2AR is expressed on fast-spiking parvalbumin-expressing (PV+) interneurons in CA1, the agonist DOI increases the firing of these cells, and restoring the receptor in PV+ interneurons alone reinstates DOI-induced anxiolysis in 5-HT2AR-null mice [11]. These cells are the main source of fast perisomatic inhibition in the hippocampus [12–14], so a sustained change in their output, as may occur with specific 5-HT2AR single nucleotide polymorphisms [15], would be expected to shift the balance of excitation and inhibition across the circuit.

PV expression is itself activity-dependent and is widely used as a proxy for the activity state of these cells [16]. The cells are ensheathed by the perineuronal net (PNN), a condensed extracellular matrix that raises PV+ excitability by lowering membrane capacitance [17,18] and by positioning ion channels and AMPA receptors at the cell surface [19,20], and enzymatic degradation of PNNs with chondroitinase ABC reduces PV+ excitability [17,18,21] and produces anxiety-like behavior [22]. The relationship between the interneuron and its net is reciprocal, since PV+ activity drives deposition of PNN constituents [19,23] while genetic loss of PNN components reduces PV+ marker expression [24,25]. Fast-spiking PV+ interneurons also carry an unusually high oxidative metabolic demand and a correspondingly elevated mitochondrial and respiratory chain content [26–28], so the proteins that supply their energy offer a further readout of their functional state.

Because 5-HT2AR activation excites the neurons that express it [4] and parvalbumin expression tracks interneuron activity [16], loss of the receptor could be expected to reduce rather than increase hippocampal PV. Whether 5-HT2AR signaling is important to PV expression and PV neuron activity, and whether any effect relies on PNN dependent or PNN independent mechanisms, remains understudied. A constitutive knockout is well suited to this question because it removes the 5-HT2AR across both development and adulthood, the periods during which PNNs are assembled and PV interneurons mature [29]. We used the BALB/c background, which displays elevated baseline anxiety relative to other inbred strains [30], because the consequences of removing a modulatory input may depend on the baseline state of the circuit it modulates. We first asked whether constitutive 5-HT2AR deletion alters anxiety-like behavior in male and female mice, using the elevated zero maze, open field, marble burying and sucrose splash tests. We then used unbiased proteomics to ask whether hippocampal PV and PNN protein levels were altered, and what else changed alongside them. Finally, to explore whether short-term 5-HT2A receptor dependent changes might occur in protein pathways that were altered in the constitutive knockouts, we also performed proteomic analyses of hippocampal tissue from wild type mice 7 days following a single dose of the selective 5-HT2AR agonist 25CN-NBOH [31].

## Materials and Methods

### Ethics statement

This study was carried out in strict accordance with the recommendations in the Guide for the Care and Use of Laboratory Animals of the National Institutes of Health. The protocol was approved by the Institutional Animal Care and Use Committee of Georgetown University (protocol number 2023-0025). Mice were anesthetized with isoflurane and euthanized by transcardial perfusion with cold PBS, and all efforts were made to minimize suffering.

### Animals

Male and female 5-HT2AR wildtype (WT) and knockout (KO) mice on the BALB/c background were used. Mice constitutively lacking the 5-HT2AR on the BALB/c background were provided by Dr. Charles D. Nichols (LSU Health Sciences Center). All animals were offspring of heterozygote breeding pairs, group-housed, and maintained under standard conditions (22 ± 2°C, 40 to 60% humidity, 12-hour light/dark cycle, lights off at 6:00 PM) with ad libitum access to standard chow (LabDiet 5001). Behavioral testing commenced at 16 weeks of age and spanned two weeks, and animals were euthanized at 18 weeks of age for tissue collection.

### Genotyping

Tail snips were collected and incubated in 200 μL GNT-K buffer (1% gelatin, 4.5% Igepal CA-630, 0.45% Tween-20, proteinase K 25 mg/mL, in Tris-KCl-Mg buffer containing 50 mM KCl, 1.5 mM MgCl2, and 10 mM Tris, pH 8.5) at 55°C for 2 hours, followed by proteinase K inactivation at 95°C for 15 minutes. PCR reactions were prepared using 12.5 μL GreenTaq, 10 μL primer stock, and 3 μL tail snip DNA. The following primers were used: HTR2A forward (5’-AAAGTCACGCTTGGCTAGAATGC-3’), HTR2A reverse (5’-GTGATGGCTCTTGATTATGCCTC-3’), and PGK (5’-AGCAGCCTCTGTTCCACATACAC-3’). Thermocycling consisted of an initial denaturation at 95°C for 1 minute, followed by touchdown cycling with annealing temperatures decreasing from 64°C to 55°C across four successive cycling blocks of 2, 6, 20, and 20 cycles respectively, with a final extension at 72°C for 10 minutes. PCR products were resolved on a 2% agarose gel. Expected band sizes were 489 bp for WT and 311 bp for KO. Heterozygous animals were identified by the presence of both bands.

### Behavioral testing

All behavioral tests were conducted between 10:00 AM and 2:00 PM during the light phase. Mice were habituated to the testing room for at least one hour before each test. Room illumination was maintained at approximately 600 lux for all tests. All apparatus were cleaned with 70% ethanol between subjects. Videos were analyzed using ANY-maze software (Stoelting, Wood Dale, IL); the marble burying and sucrose splash tests were scored by trained observers blind to genotype. Behavioral testing was conducted in both male and female animals. Tests were administered in the following order: elevated zero maze, open field test, marble burying test, and sucrose splash test, with two days between successive tests. Not all animals completed every behavioral paradigm due to equipment availability across cohorts; sample sizes for individual tests are reported in the corresponding figure captions.

For the elevated zero maze, a circular track of black acrylic (5 cm wide, 60 cm diameter, raised 50 cm) was divided into four equal quadrants, two of which were enclosed by 20 cm high walls. Each mouse was placed in the center of a closed quadrant and recorded for 5 minutes. Time in the open and closed quadrants served as the primary anxiety measures.

For the open field test, a square arena (60 × 60 × 40 cm) under even illumination was used. The center zone comprised 25% of the total area. Mice were placed in the center and recorded for 5 minutes.

For the marble burying test, mice were placed in a cage with 4.5 cm of fresh bedding and 10 glass marbles arranged in a 5 × 2 array with 2 cm gaps from the cage walls. After 30 minutes, buried marbles were counted by a trained observer blind to genotype. A marble was considered buried if at least two-thirds of its surface was covered.

For the sucrose splash test, a 30% sucrose solution was sprayed onto the dorsal coat of each mouse in its home cage. Grooming frequency and duration were recorded for 5 minutes.

Grooming included snout grooming, head washing, and body grooming. Sessions were scored in real time by a trained observer blind to genotype.

### Proteomics

Hippocampi were microdissected and protein concentration was determined by BCA assay. Samples were resuspended in 5% SDS buffer containing 50 mM TEABC and 20 mM DTT and heated at 95°C for 10 minutes. Following cooling, iodoacetamide was added to 40 mM and samples were incubated in the dark for 30 minutes. Undissolved matter was removed by centrifugation at 13,000 × g for 8 minutes and the supernatant was processed using an S-Trap column (ProtiFi, LLC). Proteins were digested with Lys-C/trypsin (Promega) overnight at 37°C and peptides were eluted and dried by SpeedVac (Fisher Scientific). Peptides were analyzed using a nanoAcquity UPLC (Waters) coupled to an Orbitrap Fusion Lumos mass spectrometer (Thermo Fisher). Samples were loaded onto a C18 trap column and separated on a BEH C18 column (75 μm × 150 mm, 1.7 μm) at 400 nL/min using a 150-minute acetonitrile gradient in data-independent acquisition (DIA) mode.

DIA raw files were analyzed using Spectronaut v17 in DirectDIA mode with default settings. A Q-value cutoff of 1% was applied at the peptide precursor and protein levels, with MS2-based quantification and local cross-run normalization. Common contaminant proteins, including keratins and immunoglobulins, were excluded prior to analysis. Differential expression between genotypes was assessed in Spectronaut, with p values adjusted for multiple testing by the Benjamini-Hochberg procedure. Proteins were considered significantly differentially expressed if they met both a false discovery rate (FDR) < 0.05 and a minimum fold change of 1.5. One sample exhibited reduced proteome coverage relative to the others; proteins not quantified in that sample were excluded from the corresponding comparisons. All proteomic analyses were conducted in male animals.

A separate cohort was used to examine the effect of 5-HT2AR activation. Wild type female BALB/c mice aged 17 weeks (n = 4 per group) received a single intraperitoneal injection of the selective 5-HT2AR agonist 25CN-NBOH (1 mg/kg) or saline vehicle and were euthanized one week later. Hippocampal tissue was collected, processed and analyzed by DIA proteomics exactly as described above, and protein sets were compared by the same rank-based method. Normalized protein group quantities for every individual animal in both cohorts are available from Dryad (doi:10.5061/dryad.xd2547f06).

### Gene Ontology enrichment analysis

Gene Ontology biological process enrichment analysis was performed using the PANTHER classification system (release 20240807) via pantherdb.org. Significantly downregulated proteins were tested against the full set of quantified proteins as the reference background rather than against the whole Mus musculus genome. Overrepresentation was assessed using Fisher’s exact test with FDR correction, and terms with FDR < 0.05 were considered significantly enriched. In a separate analysis that did not depend on the significance threshold applied to individual proteins, quantified proteins were assigned to functional sets by gene symbol and the distribution of log2 ratios within each set was compared with that of all remaining quantified proteins by two-tailed Mann-Whitney test. This rank-based comparison uses the remainder of the quantified proteome as an internal reference and is therefore insensitive to any overall difference in scaling between genotypes. Set membership is listed in Table A in S1 File.

### Statistical analysis

Unless stated otherwise, data are expressed as mean ± SEM. For behavioral measures collected in both sexes, two-way ANOVA was first conducted to assess main effects of sex, genotype, and their interaction. Where no significant effect of sex or sex by genotype interaction was detected, data were combined across sexes and compared using unpaired t-tests with Welch’s correction, which does not assume equal variances. All t-tests were two-tailed and between-subjects. Marble burying counts were bounded by the number of marbles provided and were not normally distributed, and were therefore compared by two-tailed Mann-Whitney test, with medians and interquartile ranges reported in place of mean ± SEM. Two open field recordings from KO animals were excluded prior to analysis because automated tracking failed. The threshold for significance was set at α = 0.05. Analyses were performed using GraphPad Prism 11 (GraphPad Software, Boston, MA).

## Results

### 5-HT2AR KO mice exhibit anxiety-like behavior

To assess the behavioral consequences of constitutive 5-HT2AR deletion, male and female KO and WT littermate controls on the BALB/c background were evaluated across a battery of anxiety-related behavioral paradigms (Fig 1A). In the elevated zero maze (EZM), KO animals of both sexes spent significantly less time in the open quadrants (WT: 55.41 ± 7.363 s, KO: 29.75 ± 5.312 s; t(30.92) = 2.826, p = 0.0082, n = 18 WT, 18 KO; Fig 1B) and significantly more time in the closed quadrants (WT: 244.9 ± 7.315 s, KO: 270.6 ± 5.287 s; t(30.9) = 2.846, p = 0.0078, n = 18 WT, 18 KO; Fig 1C) compared with WT controls, indicating heightened anxiety-like avoidance. In the open field test (OFT), KO animals showed a significant reduction in center zone time (WT: 42.50 ± 8.594 s, KO: 14.04 ± 4.957 s; t(20.42) = 2.868, p = 0.0094, n = 14 WT, 12 KO; Fig 1F) and a significant increase in periphery time (WT: 257.7 ± 8.585 s, KO: 286.0 ± 4.957 s; t(20.43) = 2.85, p = 0.0097, n = 14 WT, 12 KO; Fig 1G) relative to WT controls. Two KO recordings were excluded from the OFT because automated tracking failed. Together, these findings demonstrate anxiety-like behavior in 5-HT2AR KO animals across two independent paradigms. Representative open field track plots are shown in Fig 1D and 1E.

**Fig 1.**
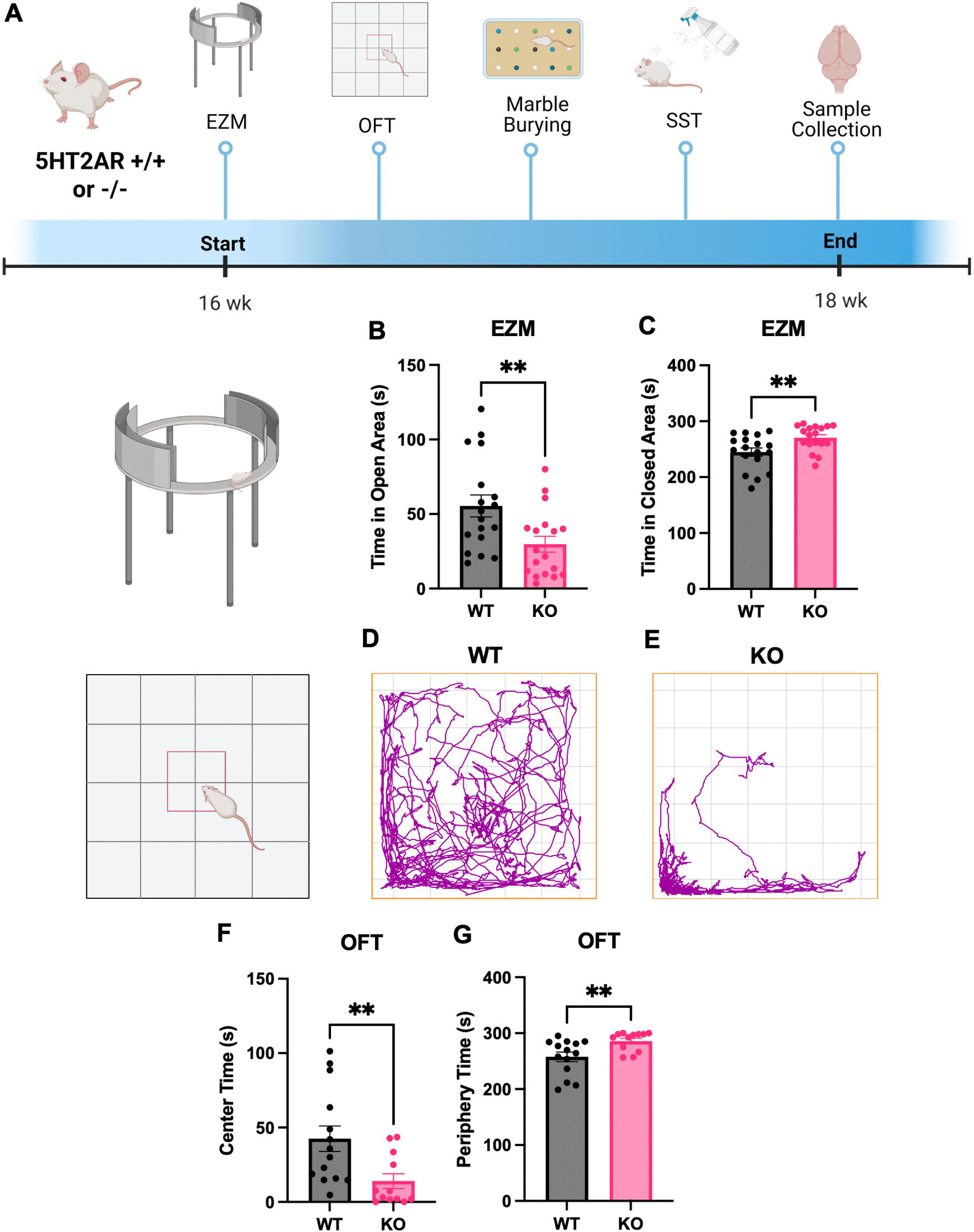
Experimental timeline and anxiety-like behavior in 5-HT2AR KO mice. (A) Schematic of the experimental timeline. Created in BioRender. Conant, K. (2027) https://BioRender.com/4i2gszb. Behavioral testing commenced at 16 weeks of age and continued through 18 weeks, at which point mice were euthanized for tissue collection and subsequent molecular analyses. (B) Time spent in open quadrants of the EZM (WT: 55.41 ± 7.363 s, KO: 29.75 ± 5.312 s; t(30.92) = 2.826, p = 0.0082, n = 18 WT, 18 KO). (C) Time spent in closed quadrants of the EZM (WT: 244.9 ± 7.315 s, KO: 270.6 ± 5.287 s; t(30.9) = 2.846, p = 0.0078, n = 18 WT, 18 KO). (D, E) Representative OFT track plots from a WT (D) and KO (E) animal. (F) Center zone time in the OFT (WT: 42.50 ± 8.594 s, KO: 14.04 ± 4.957 s; t(20.42) = 2.868, p = 0.0094, n = 14 WT, 12 KO). (G) Periphery time in the OFT (WT: 257.7 ± 8.585 s, KO: 286.0 ± 4.957 s; t(20.43) = 2.85, p = 0.0097, n = 14 WT, 12 KO). Two KO recordings were excluded because automated tracking failed. Data were combined across sexes following confirmation of no significant sex effect or sex by genotype interaction by two-way ANOVA. Unpaired t-tests with Welch’s correction. **p < 0.01. EZM, elevated zero maze; KO, knockout; OFT, open field test; WT, wildtype.

### 5-HT2AR KO mice display repetitive behavior

To further characterize the behavioral profile of 5-HT2AR KO animals, compulsive-like behavior was assessed using the marble burying test (MBT) and sucrose splash test (SST). KO animals buried significantly more marbles than WT controls (WT median 0.5, interquartile range 0 to 1.25, range 0 to 2; KO median 5.0, interquartile range 2.5 to 6.5, range 0 to 10; Mann-Whitney U = 20.50, p = 0.0001, n = 14 WT, 14 KO; Fig 2D), consistent with increased repetitive and compulsive behavior. Marble counts were compared by Mann-Whitney test rather than by t-test because burying was near the lower bound of the measure in WT animals, seven of which buried no marbles. In the SST, while this paradigm is classically employed as a measure of depressive-like behavior in terms of reduced self-grooming motivation, the phenotype observed in KO animals was inconsistent with a depressive profile and instead reflected a compulsive or perseverative grooming pattern that could reflect anxiety-driven behavior. KO animals exhibited significantly increased total grooming time (WT: 18.25 ± 2.621 s, KO: 65.53 ± 8.587 s; t(13.03) = 5.266, p < 0.001, n = 12 WT, 12 KO; Fig 2E), longer average grooming episode duration (WT: 1.045 ± 0.1098 s, KO: 2.000 ± 0.2371 s; t(15.45) = 3.653, p = 0.0023, n = 11 WT, 12 KO; Fig 2F), reduced latency to initiate grooming (WT: 24.32 ± 6.156 s, KO: 6.992 ± 1.755 s; t(11.62) = 2.707, p = 0.0195, n = 11 WT, 12 KO; Fig 2G), and a greater number of grooming episodes (WT: 17.25 ± 2.019, KO: 34.17 ± 3.047; t(19.10) = 4.628, p < 0.001, n = 12 WT, 12 KO; Fig 2H) relative to WT controls. One WT male was excluded from the average episode duration and latency measures as no grooming was observed, precluding calculation of these metrics. For latency this exclusion is conservative, since the excluded animal would have contributed the maximum possible value and its removal reduces rather than inflates the genotype difference. Together, these findings indicate that loss of 5-HT2AR signaling produces a repetitive behavioral profile consistent with heightened anxiety. Representative images of the marble array before testing and of burying in each genotype are shown in Fig 2A to 2C.

**Fig 2.**
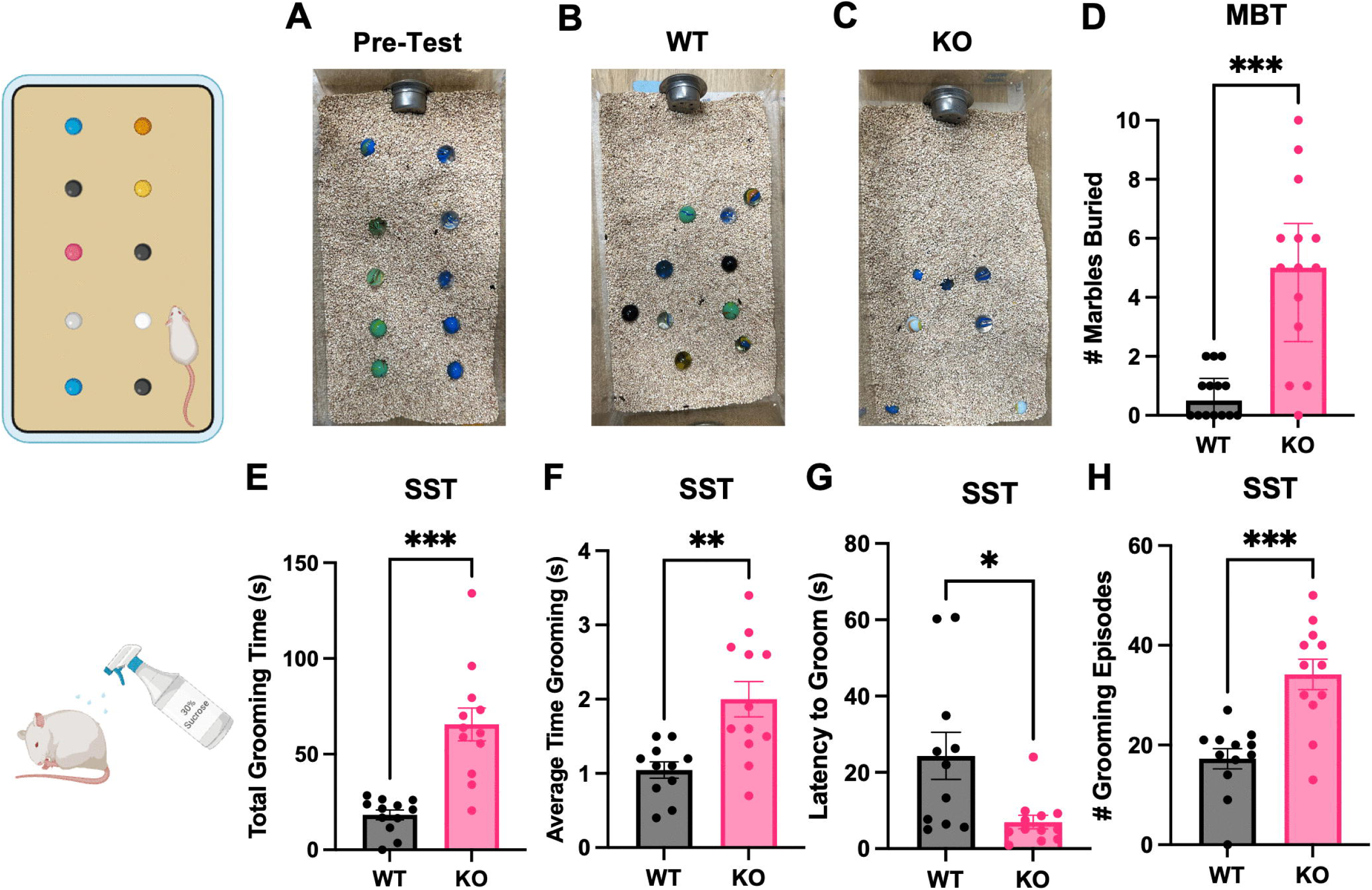
5-HT2AR KO mice display repetitive behavior. Schematic elements were created in BioRender. Conant, K. (2027) https://BioRender.com/4i2gszb. (A) Representative image of marble arrangement prior to testing. (B, C) Representative images of marble burying in WT (B) and KO (C) animals following the 30-minute testing period. (D) Number of marbles buried, shown as median with interquartile range (WT median 0.5 [0 to 1.25], KO median 5.0 [2.5 to 6.5]; Mann-Whitney U = 20.50, p = 0.0001, n = 14 WT, 14 KO). (E) Total grooming time in the SST (WT: 18.25 ± 2.621 s, KO: 65.53 ± 8.587 s; t(13.03) = 5.266, p < 0.001, n = 12 WT, 12 KO). (F) Average grooming episode duration (WT: 1.045 ± 0.1098 s, KO: 2.000 ± 0.2371 s; t(15.45) = 3.653, p = 0.0023, n = 11 WT, 12 KO). (G) Latency to initiate grooming (WT: 24.32 ± 6.156 s, KO: 6.992 ± 1.755 s; t(11.62) = 2.707, p = 0.0195, n = 11 WT, 12 KO). (H) Number of grooming episodes (WT: 17.25 ± 2.019, KO: 34.17 ± 3.047; t(19.10) = 4.628, p < 0.001, n = 12 WT, 12 KO). One WT male was excluded from panels F and G as no grooming was observed. Data were combined across sexes following confirmation of no significant sex effect or sex by genotype interaction by two-way ANOVA. Panel D, Mann-Whitney test; panels E to H, unpaired t-tests with Welch’s correction. *p < 0.05, **p < 0.01, ***p < 0.001. KO, knockout; MBT, marble burying test; SST, sucrose splash test; WT, wildtype.

### 5-HT2AR deletion is associated with reduced hippocampal PV and reduced HAPLN4 without detectable changes in aggrecan or HAPLN1

Having established a behavioral phenotype, we next asked whether hippocampal PV and PNN protein levels were altered in KO animals. Unbiased proteomic analysis of male KO and WT hippocampus quantified 4,592 protein groups following removal of common contaminants, of which 97 were significantly downregulated and 31 were significantly upregulated in KO animals (FDR < 0.05, fold change > 1.5). PV (PVALB) was among the significantly downregulated proteins, with normalized intensity reduced by approximately 54% in KO animals (WT: 346,598 ± 62,081, KO: 159,625 ± 17,904; 2.17-fold reduction, FDR = 0.0145, differential expression analysis). The corresponding unadjusted comparison of normalized intensities is shown in Fig 3A (t(4.661) = 2.894, p = 0.0370) and is descriptive of the same data rather than an independent test. In contrast, two core PNN structural components, hyaluronan and proteoglycan link protein 1 (HAPLN1) and aggrecan (ACAN), did not differ significantly between genotypes (Fig 3B and 3C; HAPLN1, WT: 793,857 ± 134,064, KO: 672,529 ± 68,122; t(5.936) = 0.8068, p = 0.4509, n = 5 per group. ACAN, WT: 44,494 ± 4,541, KO: 39,372 ± 2,555; t(4.837) = 0.9830, p = 0.3722, n = 4 WT, 5 KO). ACAN was not detected in one WT sample, which was therefore excluded from that comparison. A third matrix protein behaved differently. Hyaluronan and proteoglycan link protein 4 (HAPLN4) was among the significantly downregulated proteins, with normalized intensity reduced by approximately 79% in KO animals (WT: 113,367 ± 22,242, KO: 23,752 ± 6,607; 4.77-fold reduction, FDR = 0.0073, differential expression analysis). The corresponding unadjusted comparison of normalized intensities is shown in Fig 3D (t(4.701) = 3.862, p = 0.0133). The reduction in PV was therefore accompanied by a marked reduction in HAPLN4 but not by detectable changes in ACAN or HAPLN1 (Fig 3). With n = 5 per group and a pooled coefficient of variation of 32% for HAPLN1 and 18% for ACAN, these comparisons had approximately 25% and 57% power respectively to detect a 30% difference, so the absence of a significant change should not be read as evidence that the matrix is unaffected.

**Fig 3.**
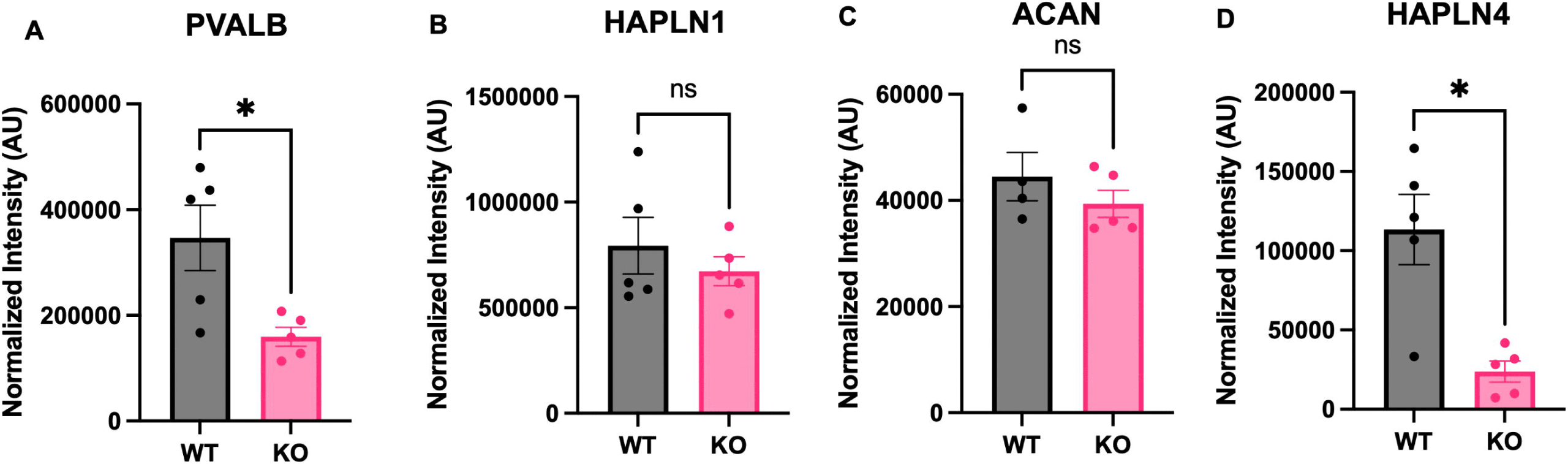
5-HT2AR deletion is associated with reduced hippocampal PV and reduced HAPLN4 without detectable changes in aggrecan or HAPLN1. (A) Normalized PVALB intensity (WT: 346,598 ± 62,081, KO: 159,625 ± 17,904; t(4.661) = 2.894, p = 0.0370, n = 5 per group). (B) Normalized HAPLN1 intensity (WT: 793,857 ± 134,064, KO: 672,529 ± 68,122; t(5.936) = 0.8068, p = 0.4509, n = 5 per group). (C) Normalized ACAN intensity (WT: 44,494 ± 4,541, KO: 39,372 ± 2,555; t(4.837) = 0.9830, p = 0.3722, n = 4 WT, 5 KO). ACAN was not detected in one WT sample, which was excluded from that comparison. (D) Normalized HAPLN4 intensity (WT: 113,367 ± 22,242, KO: 23,752 ± 6,607; t(4.701) = 3.862, p = 0.0133, n = 5 per group). All four panels display normalized intensities from the same proteomic dataset; the plotted t-tests are descriptive of the differential expression analysis and are not independent tests. All analyses were conducted in male hippocampal tissue. Unpaired t-tests with Welch’s correction. *p < 0.05; ns, not significant. ACAN, aggrecan; HAPLN1, hyaluronan and proteoglycan link protein 1; HAPLN4, hyaluronan and proteoglycan link protein 4; KO, knockout; PV, parvalbumin; PVALB, parvalbumin alpha; WT, wildtype.

### Energy metabolism and translational machinery proteins are reduced in 5-HT2AR KO hippocampus

To determine whether the downregulated proteins were enriched for particular biological processes beyond those anticipated by the PV hypothesis, Gene Ontology biological process enrichment analysis was performed on the 97 downregulated protein groups (Fig 4A), which carried 99 gene identifiers of which 94 mapped to the PANTHER database, using the 4,624 mapped identifiers from the quantified proteome as the reference background. Eight terms reached significance, all relating to mitochondrial respiration (Fig 4B). Oxidative phosphorylation showed the strongest statistical enrichment (12 proteins observed, 1.77 expected; 6.79-fold; FDR = 1.33 × 10□³), alongside mitochondrial ATP synthesis coupled electron transport (9 proteins; 8.68-fold; FDR = 2.17 × 10□³), ATP synthesis coupled electron transport (9 proteins; 8.51-fold; FDR = 1.94 × 10□³), aerobic electron transport chain (8 proteins; 8.56-fold; FDR = 5.35 × 10□³), respiratory electron transport chain (10 proteins; 7.57-fold; FDR = 2.84 × 10□³), electron transport chain (10 proteins; 7.13-fold; FDR = 2.02 × 10□³), aerobic respiration (13 proteins; 4.54-fold; FDR = 6.29 × 10□³), and cellular respiration (13 proteins; 4.18-fold; FDR = 1.36 × 10□²). Enrichment analysis of the 31 upregulated proteins returned no significantly enriched terms. These terms are hierarchically related within the Gene Ontology and were driven by a shared set of 12 proteins spanning all four respiratory complexes, including subunits of complex I (NDUFS2, NDUFS4, NDUFS5, NDUFA7, NDUFA8), complex II (SDHD), complex III (UQCRH, UQCRFS1), and complex IV (COX6C, COX7A2L), together with cytochrome c (CYCS) and the ubiquinone biosynthesis enzyme COQ7 (Fig 4C). Reductions distributed across the entire respiratory chain, rather than confined to a single complex, are more consistent with a general decrease in respiratory chain protein abundance than with a defect in the assembly of an individual complex.

**Fig 4.**
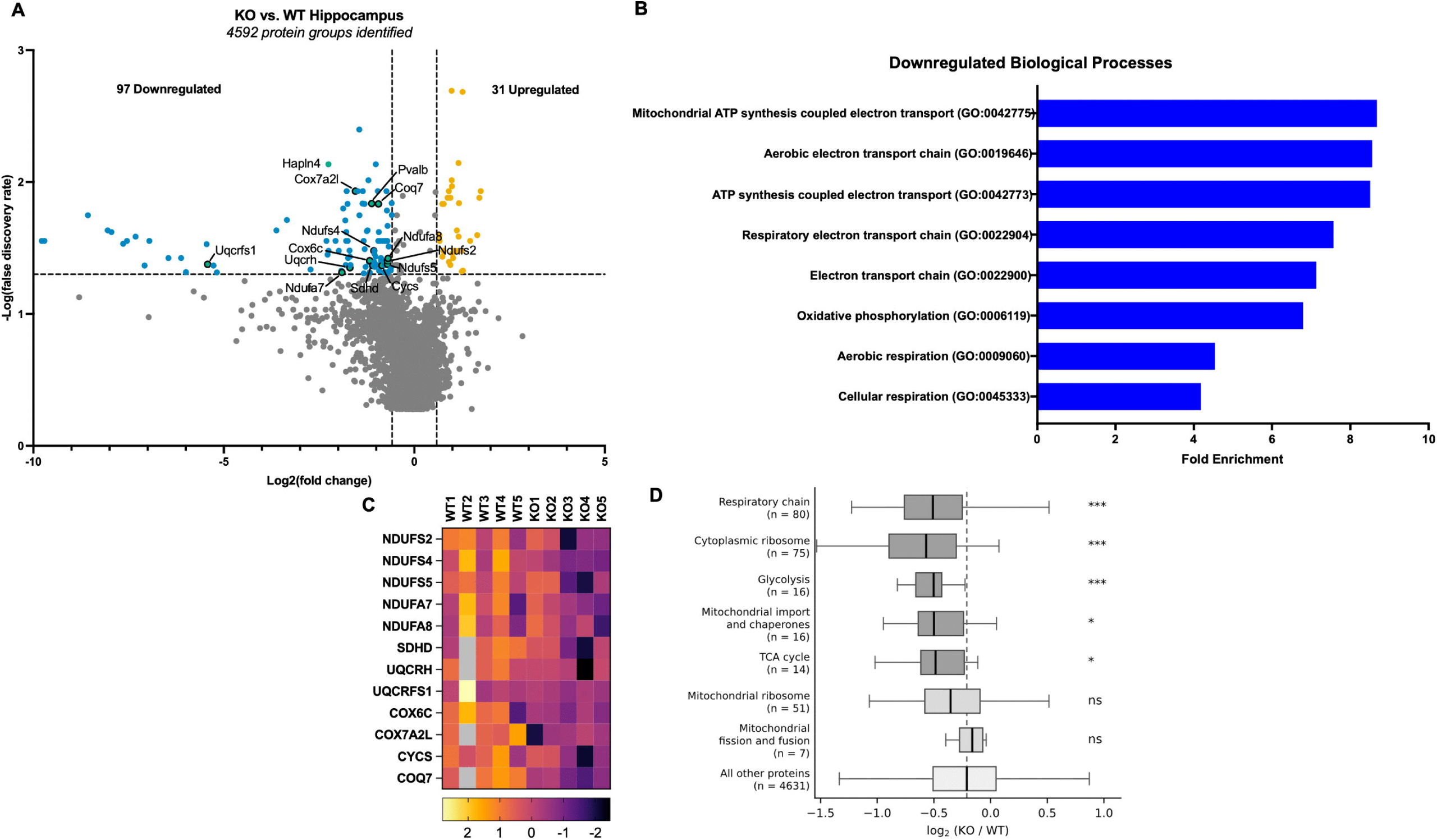
Energy metabolism and translational machinery proteins are reduced in 5-HT2AR KO hippocampus. (A) Volcano plot of 4,592 protein groups quantified by DIA proteomic analysis of male KO and WT hippocampus. Dotted lines indicate thresholds of FDR < 0.05 and fold change > 1.5. Downregulated proteins are shown in blue (n = 97), upregulated proteins in orange (n = 31), and non-significant proteins in grey. The 12 respiratory chain proteins driving the enrichment shown in panel B are highlighted in green. (B) Gene Ontology biological process enrichment analysis of downregulated proteins, plotted as fold enrichment. Eight terms reached significance (FDR < 0.05), all relating to mitochondrial respiration. Enrichment was calculated using the quantified proteome as the reference background (94 of the 99 gene identifiers carried by the 97 downregulated protein groups, and 4,624 of 4,631 identifiers from the quantified proteome, mapped to the PANTHER database). Enriched terms are hierarchically related within the Gene Ontology and reflect an overlapping set of proteins. (C) Heat map of the 12 respiratory proteins driving the enrichment shown in panel B, displayed as row z-scores of log2-transformed normalized intensities across individual samples. Proteins include subunits of complex I (NDUFS2, NDUFS4, NDUFS5, NDUFA7, NDUFA8), complex II (SDHD), complex III (UQCRH, UQCRFS1), and complex IV (COX6C, COX7A2L), together with cytochrome c (CYCS) and the ubiquinone biosynthesis enzyme COQ7. Grey indicates that a protein was not quantified in that sample. (D) Distribution of log2 ratios for defined protein sets compared with all other quantified proteins. Boxes show the median and interquartile range, whiskers extend to 1.5 times the interquartile range, and the dashed line marks the median of the quantified proteome. Each set was compared with all remaining quantified proteins by two-tailed Mann-Whitney test. Set membership is listed in Table S1. *p < 0.05, ***p < 0.001; ns, not significant. All analyses were conducted in male hippocampal tissue, n = 5 per group. DIA, data-independent acquisition; FDR, false discovery rate; KO, knockout; WT, wildtype.

Because enrichment analysis evaluates only the proteins that cross a significance threshold, we next asked how broadly energy metabolism was affected by comparing whole protein sets against the rest of the quantified proteome. Cytoplasmic ribosomal proteins showed the largest coordinated shift (n = 75, median log2 ratio −0.569 compared with −0.203 for all other proteins, p = 6.0 × 10□¹²). Respiratory chain proteins were similarly shifted downward (n = 80, −0.509, p = 1.3 × 10□□), as were enzymes of glycolysis (n = 16, −0.503, p = 5.6 × 10□□), the tricarboxylic acid cycle (n = 14, −0.486, p = 0.037), and the mitochondrial protein import and chaperone machinery (n = 16, −0.500, p = 0.015). In contrast, mitochondrial ribosomal proteins (n = 51, −0.353, p = 0.154) and proteins governing mitochondrial fission and fusion (n = 7, −0.162, p = 0.77) did not differ from the rest of the proteome. The reduction therefore extended to glycolytic as well as oxidative energy-producing enzymes and to the cytoplasmic translational machinery, while the machinery of mitochondrial gene expression and mitochondrial dynamics was comparatively spared. This pattern is more consistent with a coordinated reduction in biosynthetic and bioenergetic capacity than with a reduction in mitochondrial number (Fig 4D).

### A single dose of the 5-HT2AR agonist 25CN-NBOH shifts the same protein sets in the opposite direction

The reductions described above followed loss of the 5-HT2AR across the lifespan. To ask whether 5-HT2AR activation moves the same protein sets, hippocampal tissue was examined from a separate cohort of wild type female BALB/c mice given a single intraperitoneal dose of the selective 5-HT2AR agonist 25CN-NBOH (1 mg/kg) or saline vehicle and sampled one week later (n = 4 per group). Applying the same set-level comparison, respiratory chain proteins were shifted upward relative to the remainder of the proteome (set median +0.060, p = 0.0023), as were tricarboxylic acid cycle enzymes (+0.158, p = 0.036) and cytoplasmic ribosomal proteins (+0.054, p = 0.024), in each case opposite in direction to the 5-HT2AR knockout (Fig S1). The magnitude of these shifts was approximately one fifth of that seen after 5-HT2AR deletion. In contrast, PVALB (log2 ratio +0.287, FDR = 0.354), HAPLN4 (+0.192, FDR = 0.157) and the remaining matrix constituents did not differ between agonist and vehicle groups.

## Discussion

Constitutive deletion of the 5-HT2AR produced anxiety-like and repetitive behavior across four paradigms in both sexes, and hippocampal tissue from these animals showed reduced parvalbumin and reduced HAPLN4 alongside coordinated reductions in the proteins that supply energy and support protein synthesis, with aggrecan and HAPLN1 unchanged. Recent work has shown that 5-HT2AR is expressed on fast-spiking PV+ interneurons in CA1, that the agonist DOI increases their firing, and that restoring the 5-HT2AR in PV+ interneurons alone reinstates DOI-induced anxiolysis in 5-HT2AR-null mice [11]. The present findings indicate that the 5-HT2AR also shapes the baseline characteristics of these cells.

Loss of the 5-HT2AR increased anxiety-like behavior on the standard open area measures in both the elevated zero maze and the open field [32–35], which places endogenous 5-HT2AR signaling on the anxiolytic side and is consistent with the acute anxiolytic action of 5-HT2AR agonists [11]. Reduced grooming in the sucrose splash test is normally taken as evidence of diminished self-care and a depressive-like state [36]. However, the result runs opposite to the conventional reading of the assay. 5-HT2AR knockout animals groomed more, for longer, more often, and sooner. Read alongside the increase in marble burying [37], the pattern is better described as perseverative than as depressive [38,39]. One prior study appears to contradict the anxiety findings. Mice lacking the 5-HT2AR on a 129S6/SvEv background show reduced conflict anxiety, restored by cortical expression of the 5-HT2AR [9]. BALB/c and 129S6/SvEv differ in baseline anxiety [30], and the two studies also differ in the behavioral batteries used and in the ages at testing. Because BALB/c animals are already anxious at baseline, removing an anxiolytic input has more room to act.

Acute agonism of the 5-HT2AR reduces anxiety by acting on hippocampal PV interneurons [11]. PV expression is regulated by the activity of the interneurons that express it and is widely used as a proxy for that activity [16]. That activity is in turn supported by the perineuronal net, which raises excitability by lowering membrane capacitance and by concentrating ion channels and AMPA receptors at the cell surface [17–20]. The 5-HT2AR is excitatory [4], so its absence may remove a depolarizing input to these cells. Reduced PV in 5-HT2AR knockout hippocampus could therefore follow from reduced PV interneuron activity. The matrix itself, however, was largely intact. HAPLN1, which is critical for net assembly [29,40], and aggrecan, which is abundant in hippocampal nets and increases with PV activity [40,41], did not differ between genotypes. The link protein HAPLN4 was, however, reduced. That dissociation is informative, because PV activity normally drives deposition of matrix constituents [19,23] and genetic loss of those constituents reduces PV marker expression [24,25], so the two ordinarily move together. HAPLN4, however, merits particular attention. Link proteins stabilize the attachment of chondroitin sulfate proteoglycans to the hyaluronan backbone, and in their absence the net fails to condense [29]. HAPLN4 binds hyaluronan and colocalizes largely with brevican [42], and mice lacking it show altered extracellular space volume and diffusion properties in the thalamus and in the auditory system [43,44]. Loss of this single link protein therefore has measurable consequences for the matrix even where the bulk constituents remain. Recent single cell work identifies HAPLN4 as one of the few genes that predicts both whether a PV interneuron carries a perineuronal net and how intense that net is, whereas parvalbumin itself predicts the presence of a net but not its intensity [45]. A selective reduction in HAPLN4 alongside unchanged aggrecan and HAPLN1 therefore points toward altered organization of the net rather than a loss of matrix material [45]. Intact nets have been reported alongside reduced PV expression in Shank 3 mutation autism models [46], and thus a matrix that appears unchanged in bulk need not be functionally unchanged. Direct histological confirmation will be required.

The proteomic changes point in the same direction. Proteins of the respiratory chain, the tricarboxylic acid cycle, glycolysis and the cytoplasmic ribosome were all reduced, whereas mitochondrial ribosomal proteins and the machinery of mitochondrial fission and fusion were not. Mitoribosomes translate the thirteen mitochondrially encoded subunits of the respiratory chain [47], and the fission and fusion machinery is regulated as part of mitochondrial quality control [48], so neither set is a passive index of how many mitochondria are present. A uniform loss of mitochondria would nonetheless have lowered all of these together, so the pattern is better described as selective remodeling than as a change in mitochondrial number [49]. The fall in cytoplasmic ribosomal proteins, the largest coordinated shift in the dataset, points separately to reduced capacity for protein synthesis in the cytosol. The same dissociation has been reported directly in PV interneurons after chronic fluoxetine, a treatment that also lowers PV expression and PNNs [50], where respiratory chain and ATP synthase transcripts fell while mitochondrial DNA content rose [51]. This matters for PV interneurons in particular, because fast-spiking cells carry an unusually high oxidative demand and depend on uninterrupted ATP production to restore ion gradients after each action potential [26–28], which leaves them less able than other neurons to tolerate a fall in that supply [28] and selectively vulnerable in models of oxidative stress [52,53]. The same metabolic program separates PNN-bearing from PNN-lacking PV interneurons in recent single-cell data, where it marks the fast-spiking specialized population [45], so its reduction here may reflect loss of that specialization rather than an independent mitochondrial deficit.

That capacity appears to be bidirectional. A single dose of 25CN-NBOH, which is selective for 5-HT2AR over the closely related 5-HT2CR [31], shifted the same protein sets in the opposite direction one week later, with an effect roughly one fifth the magnitude of 5-HT2AR deletion (Fig S1). Removing the receptor lowers these proteins and activating it raises them, which places 5-HT2AR signaling upstream of hippocampal metabolic capacity rather than alongside it, and shows that this capacity is still under receptor control in the adult animal. That one exposure leaves a trace seven days after the drug has cleared is consistent with evidence that a single dose of a 5-HT2AR agonist produces structural changes in cortex that persist for weeks [10]. PV, HAPLN4 and the matrix constituents did not move, and that separation assigns the two phenotypes to different time and/or developmental windows, with the parvalbumin and link protein reductions perhaps better attributed to the absence of the receptor during the postnatal period in which these interneurons mature [54,55]. Two limitations apply to the comparison. The agonist cohort was female whereas the 5-HT2AR knockout proteomics was conducted in males, so manipulation and sex are confounded, and the two datasets were acquired separately.

Although a constitutive knockout cannot separate developmental from adult contributions, the direction of the molecular phenotype is informative for the balance between excitation and inhibition in the hippocampus. PV interneurons supply the majority of perisomatic inhibition onto principal cells [12–14], and PV expression tracks the activity of those interneurons [16]. Because the 5-HT2AR supplies excitatory drive to these inhibitory cells [4], removing it may reduce inhibitory output even though the lost input was itself excitatory. A sustained reduction in PV, together with reduced HAPLN4, would be expected to lower inhibitory drive onto principal neurons and shift this balance toward excitation. Because increased excitatory drive in the hippocampus can be associated with heightened anxiety-like behavior, this offers a direct link between 5-HT2AR loss and the behavioral phenotype reported here, and it does not require the 5-HT2AR to be acting acutely at the time of testing.

Several limitations bear on interpretation. The 5-HT2AR is absent from conception and in all tissues, so this model cannot separate developmental from adult contributions, and conditional deletion will be required to do so since serotonergic signaling during early postnatal windows shapes adult emotional behavior [54,55]. Proteomic analysis was restricted to male animals and to whole hippocampus, so the molecular phenotype cannot be assumed to generalize across sexes or along the dorsoventral axis, where the ventral pole is more closely tied to anxiety-related behavior [11].

In summary, constitutive loss of the 5-HT2AR produced anxiety-like and repetitive behavior on the BALB/c background. This behavioral profile differs from that observed with the receptor knockout on the 129S6/SvEv background. In the present study we also observed reduced hippocampal PV, reduced HAPLN4, and coordinated reductions in the proteins that supply energy and support protein synthesis in the knockout animals. Aggrecan and HAPLN1 were unchanged, pointing to a change in the state of these interneurons rather than disassembly of the perineuronal net. Moreover, a single dose of a 5-HT2A selective agonist moved the metabolic proteins in the opposite direction of that observed in knockouts. Together, these data support the possibility that hippocampal PV interneurons are particularly sensitive to short and long term alterations in 5-HT2A dependent signaling.

## Acknowledgments

The authors thank Dr. Charles D. Nichols, Department of Pharmacology and Experimental Therapeutics, LSU Health Sciences Center, New Orleans, LA, for generously providing the 5-HT2AR KO mouse line used in this study. The authors also thank Dr. William C. Wetsel, Department of Psychiatry and Behavioral Sciences and Director of the Mouse Behavioral and Neuroendocrine Analysis Core Facility, Duke University Medical Center, Durham, NC, for insightful advice regarding the behavioral studies.

## Supporting information

Fig S1. Protein set changes after 5-HT2AR deletion and after agonist exposure. Distribution of log2 ratios for each protein set relative to the median of the remainder of the same proteome, for 5-HT2AR knockout versus wild type and for 25CN-NBOH versus saline. Boxes show the median and interquartile range, whiskers extend to 1.5 times the interquartile range, and the dashed line marks no change relative to the proteome median. Set sizes are given for the 5-HT2AR knockout and agonist datasets respectively. Each set was compared with all remaining quantified proteins in its own dataset by two-tailed Mann-Whitney test. *p < 0.05, **p < 0.01, ***p < 0.001; ns, not significant.

Table S1. Protein set membership and set-level comparisons. Gene symbols assigned to each functional protein set, with the number of proteins quantified, the median log2 ratio, and the Mann-Whitney p value against all remaining quantified proteins.

## Notes

https://www.doi.org/10.5061/dryad.xd2547f06

### Competing Interest Statement

The authors have declared no competing interest.

