## Supporting Information for "5-HT2A receptor deletion is associated with reduced hippocampal parvalbumin levels and anxiety-like behavior in BALB/c mice"

**Fig S1. Protein set changes after 5-HT2AR deletion and after agonist exposure.**

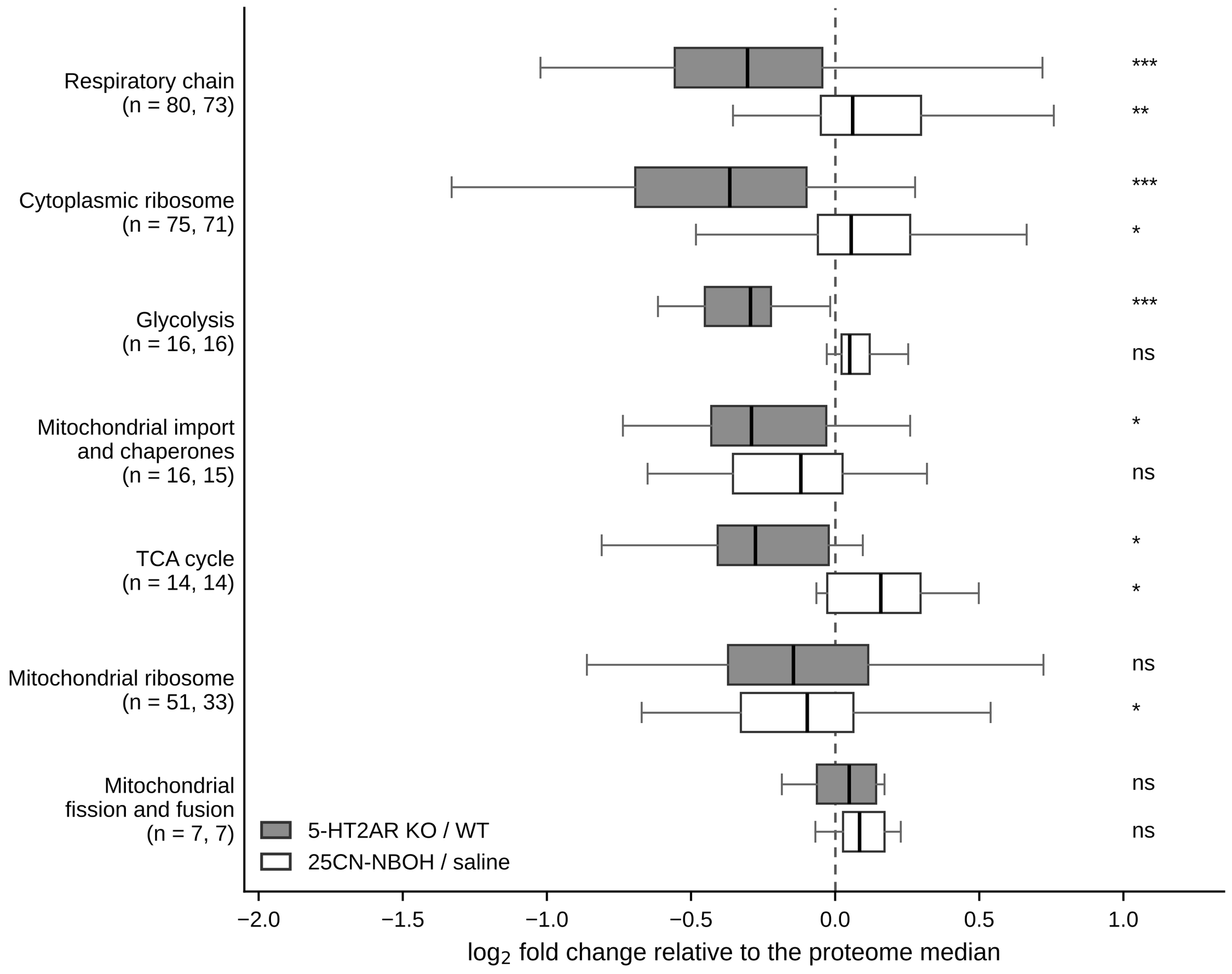

Distribution of log2 ratios for each protein set relative to the median of the remainder of the same proteome, for 5-HT2AR knockout versus wild type and for 25CN-NBOH versus saline. Boxes show the median and interquartile range, whiskers extend to 1.5 times the interquartile range, and the dashed line marks no change relative to the proteome median. Set sizes are given for the 5-HT2AR knockout and agonist datasets respectively. Each set was compared with all remaining quantified proteins in its own dataset by two-tailed Mann-Whitney test. *p < 0.05, **p < 0.01, ***p < 0.001; ns, not significant.

**Table S1. Protein set membership and set-level comparisons.**

Gene symbols assigned to each functional protein set, with the number of proteins quantified, the median log2 ratio for 5-HT2AR knockout versus wild type, and the two-tailed Mann-Whitney p value against all remaining quantified proteins in the same dataset.

| **Protein set** | **n quantified** | **Median log2 KO/WT** | **Mann-Whitney p** |
| --- | --- | --- | --- |
| Respiratory chain | 80 | -0.509 | 1.26e-07 |
| Cytoplasmic ribosome | 75 | -0.569 | 5.96e-12 |
| Glycolysis | 16 | -0.503 | 5.64e-04 |
| Mitochondrial import and chaperones | 16 | -0.5 | 1.54e-02 |
| TCA cycle | 14 | -0.486 | 3.73e-02 |
| Mitochondrial ribosome | 51 | -0.353 | 1.54e-01 |
| Mitochondrial fission and fusion | 7 | -0.162 | 7.72e-01 |

**Gene symbols by set**

**Respiratory chain (n = 80).** Atp5f1a, Atp5f1b, Atp5f1c, Atp5f1d, Atp5f1e, Atp5mc1, Atp5mc2, Atp5mc3, Atp5me, Atp5mf, Atp5mg, Atp5mk, Atp5mpl, Atp5pb, Atp5pd, Atp5pf, Atp5po, Cox15, Cox18, Cox20, Cox4i1, Cox5a, Cox5b, Cox6a1, Cox6b1, Cox6c, Cox7a2, Cox7a2l, Cycs, Ndufa10, Ndufa11, Ndufa12, Ndufa13, Ndufa2, Ndufa3, Ndufa4, Ndufa5, Ndufa6, Ndufa7, Ndufa8, Ndufa9, Ndufaf1, Ndufaf2, Ndufaf3, Ndufaf4, Ndufaf5, Ndufaf6, Ndufb1, Ndufb10, Ndufb11, Ndufb3, Ndufb4, Ndufb5, Ndufb6, Ndufb7, Ndufb8, Ndufb9, Ndufc2, Ndufs1, Ndufs2, Ndufs3, Ndufs4, Ndufs5, Ndufs6, Ndufs7, Ndufs8, Ndufv1, Ndufv2, Ndufv3, Sdha, Sdhb, Sdhc, Sdhd, Uqcr10, Uqcrb, Uqcrc1, Uqcrc2, Uqcrfs1, Uqcrh, Uqcrq

**Cytoplasmic ribosome (n = 75).** Rpl10, Rpl10a, Rpl11, Rpl12, Rpl13, Rpl13a, Rpl14, Rpl15, Rpl17, Rpl18, Rpl18a, Rpl19, Rpl21, Rpl22, Rpl22l1, Rpl23, Rpl23a, Rpl24, Rpl26, Rpl27, Rpl27a, Rpl28, Rpl29, Rpl3, Rpl30, Rpl31, Rpl32, Rpl34, Rpl35, Rpl35a, Rpl36, Rpl36a, Rpl37a, Rpl38, Rpl39, Rpl4, Rpl5, Rpl6, Rpl7, Rpl7a, Rpl8, Rpl9, Rps10, Rps11, Rps12, Rps13, Rps14, Rps15, Rps15a, Rps16, Rps17, Rps18, Rps19, Rps2, Rps20, Rps21, Rps23, Rps24, Rps25, Rps26, Rps27, Rps27a, Rps27l, Rps28, Rps29, Rps3, Rps3a, Rps4x, Rps5, Rps6, Rps6ka2, Rps6ka5, Rps7, Rps8, Rps9

**Glycolysis (n = 16).** Aldoa, Aldoc, Eno1, Eno2, Eno3, Gapdh, Hk1, Ldha, Ldhb, Pfkl, Pfkm, Pfkp, Pgam1, Pgk1, Pkm, Tpi1

**Mitochondrial import and chaperones (n = 16).** Hspa9, Hspd1, Hspe1, Timm13, Timm22, Timm23, Timm29, Timm44, Timm50, Timm9, Tomm20, Tomm22, Tomm34, Tomm40, Tomm40l, Tomm70

**TCA cycle (n = 14).** Aco2, Cs, Dld, Dlst, Idh2, Idh3a, Idh3g, Mdh2, Ogdh, Pdha1, Pdhb, Sucla2, Suclg1, Suclg2

**Mitochondrial ribosome (n = 51).** Mrpl10, Mrpl11, Mrpl12, Mrpl13, Mrpl15, Mrpl16, Mrpl17, Mrpl19, Mrpl2, Mrpl21, Mrpl22, Mrpl23, Mrpl24, Mrpl27, Mrpl28, Mrpl3, Mrpl32, Mrpl37, Mrpl38, Mrpl39, Mrpl4, Mrpl41, Mrpl44, Mrpl45, Mrpl46, Mrpl48, Mrpl49, Mrpl51, Mrpl53, Mrpl55, Mrpl58, Mrpl9, Mrps10, Mrps11, Mrps14, Mrps15, Mrps16, Mrps18a, Mrps21, Mrps22, Mrps23, Mrps26, Mrps27, Mrps30, Mrps31, Mrps34, Mrps35, Mrps36, Mrps5, Mrps7, Mrps9

**Mitochondrial fission and fusion (n = 7).** Chchd3, Chchd6, Dnm1l, Fis1, Immt, Mfn2, Opa1
